# CATFISHR: integration of transcriptome and inferred CNV profiles for comprehensive malignant cell identification

**DOI:** 10.64898/2026.10.22.760906

**Authors:** Prachi Shukla, Maria Mannino, Myriam Hofer, Evelyn Lattmann, Veronika Haunerdinger, Patrick Turko, Mitchell P. Levesque

## Abstract

Malignant cell identification in single-cell RNA sequencing (scRNA-seq) remains challenging because transcriptomic cell typing, somatic variant detection and inferred copy-number approaches each have modality-specific limitations. Rare or ambiguous malignant populations are especially vulnerable to misclassification or exclusion, despite their relevance to treatment resistance, minimal residual disease, and metastasis. We developed CATFISHR, a semi-automated workflow that integrates RNA-expression similarity, inferred copy-number variation (CNV) profiles, and consensus cell typing to classify malignant cells relative to high-confidence references. CATFISHR was evaluated across eight cohorts, including seven scRNA-seq datasets spanning multiple carcinomas and melanoma subtypes, and one 10x Visium spatial transcriptomics sample. Compared with available benchmarks or ground truths, CATFISHR achieved balanced accuracy > 0.85 in six of eight cohorts and F1 score > 0.80 in six of eight cohorts. Cells uniquely identified by CATFISHR as malignant (“Hidden” cells) were present in every dataset and were not consistently explained by standard quality-control metrics. In several cohorts, Hidden cells were transcriptionally closer to benchmark malignant than non-malignant compartments; in Visium data, sampled Hidden spots contained histologist-confirmed morphologically malignant cells. CATFISHR reduces missed malignant calls while preserving rare, transitional and benchmark-discordant populations for downstream investigation.

**GRAPHICAL ABSTRACT:** 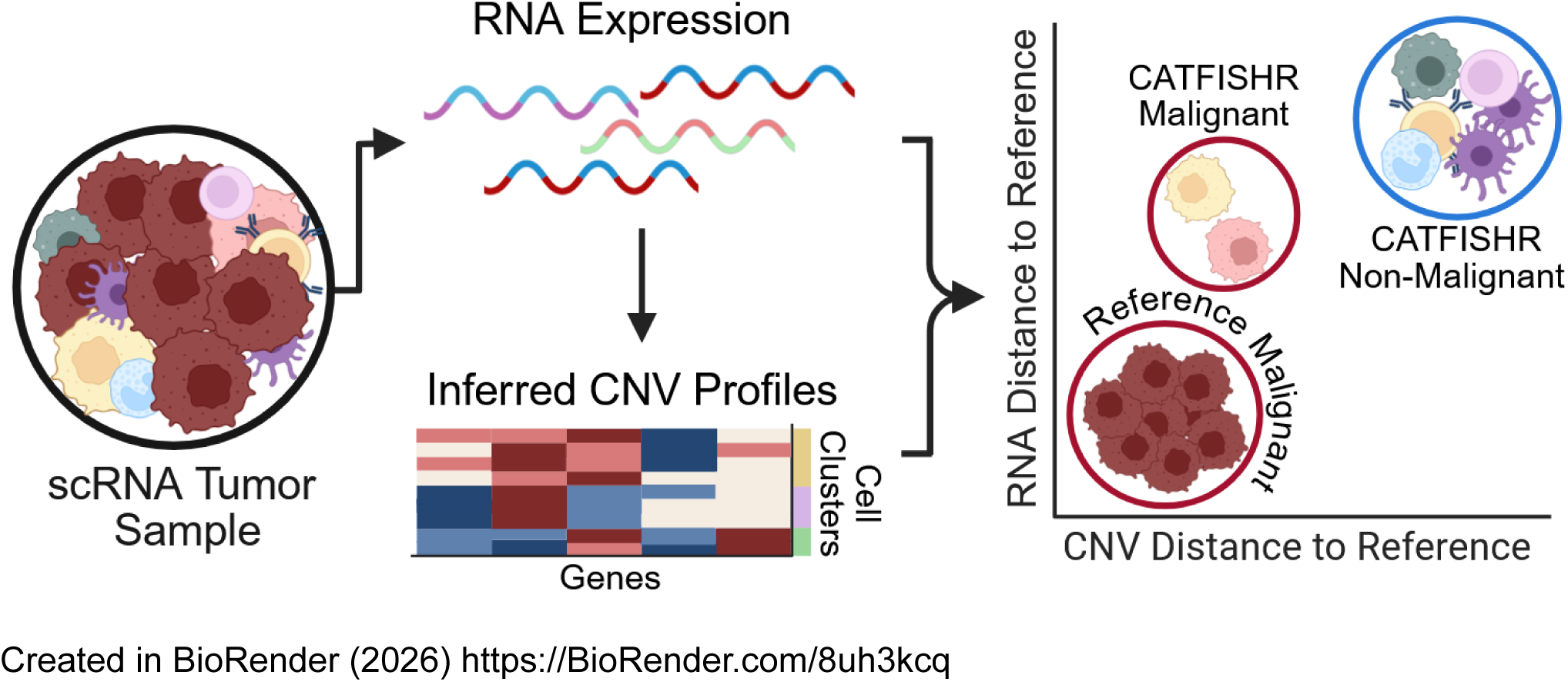

## INTRODUCTION

Accurate identification of malignant cells in single cell RNA sequencing data remains a central challenge in cancer biology. Reliable malignant cell classification is critical for downstream analyses that require separation of tumor-intrinsic signals from the tumor microenvironment, including characterization of malignant cell states, immune and stromal composition, and intratumoral transcriptional and clonal heterogeneity. Several clinically relevant populations that impact resistance, minimal residual disease, and metastasis are rare or show hybrid features, making accurate classification especially important (1–3).

Cell typing alone is often insufficient to distinguish malignant cells, as it can be confounded by tumor plasticity, the presence of non-malignant cells of the same lineage, and shared lineage programs (4). Although single-nucleotide variants (SNVs) may help distinguish malignant from non-malignant cells, they are not reliably detected at single-cell resolution in short-read 10x Chromium data (5–7).

In addition to cell typing, deletions and amplifications of genomic segments arising from genomic instability can be leveraged to identify malignant cells. These large-scale somatic copy-number alterations, hereafter referred to as CNVs, are a hallmark of cancer, with aneuploidy observed in 90% of solid tumors and 76% of hematological tumors (8). Thus, CNV profiles provide an orthogonal signal to support malignant cell identification (9, 10). Most automated tools for tumor cell identification from scRNA-seq, including InferCNV, CopyKAT (9), SCEVAN (11), and Numbat (12), focus primarily on CNV inference, with cell typing performed in parallel. However, CNV inference can be noisy in low-depth scRNA-seq and sensitive to parameter settings and reference-cell choice. Moreover, clonality, changes in phenotype, cancer-type aneuploidy/CNA burden, immune-cell-specific genomic rearrangements, such as V(D)J recombination, and technical effects can further complicate inferred CNV output and downstream malignancy assignment (13, 14).

To address these limitations, several studies identify malignant cells using multi-step, de novo workflows that apply RNA-based cell type filtering to select for the cancer cell type and then CNV-based clustering, thresholding, and/or application of gene signatures to label malignant clusters (15–18). However, sequential workflows evaluate each feature space independently, such that cells excluded at earlier steps are not assessed within subsequent feature spaces. This precludes the characterization of cells misclassified by one approach but supported by another. To our knowledge, few workflows explicitly integrate transcriptomic similarity and inferred CNV similarity from scRNA-seq within a joint malignancy-calling framework, rather than applying these evidence sources independently or sequentially (14, 18).

To address this challenge, we developed a semi-automated method to identify malignant cells called inferred CNV And Transcriptome Framework for Identification of Similarity to High-confidence References (CATFISHR). CATFISHR integrates RNA expression with inferred CNV profiles to account for cells which are ambiguous in one space but resolved in the other. An overview of the workflow is shown in Figure 1. Briefly, transcriptomic scRNA expression and inferred CNVs are analyzed in two parallel workflows, beginning with clustering in the respective RNA and CNV spaces. From these clusters, “malignant references” are selected based on agreement within cell typing algorithms (RNA), or visual malignant cluster identification from residual copy number heatmaps (CNV). The similarity of each cell to the malignant reference centroids is calculated using the Mahalanobis distance metric within each feature space. Cells with low distances to reference clusters in both RNA and CNV space are classified as malignant, indicating that their transcriptomic and copy-number profiles are concordant with the reference malignant clusters. To formalize this classification, mean shift clustering is applied in the integrated RNA–CNV space to separate malignant from non-malignant cell populations.

**Figure 1.**
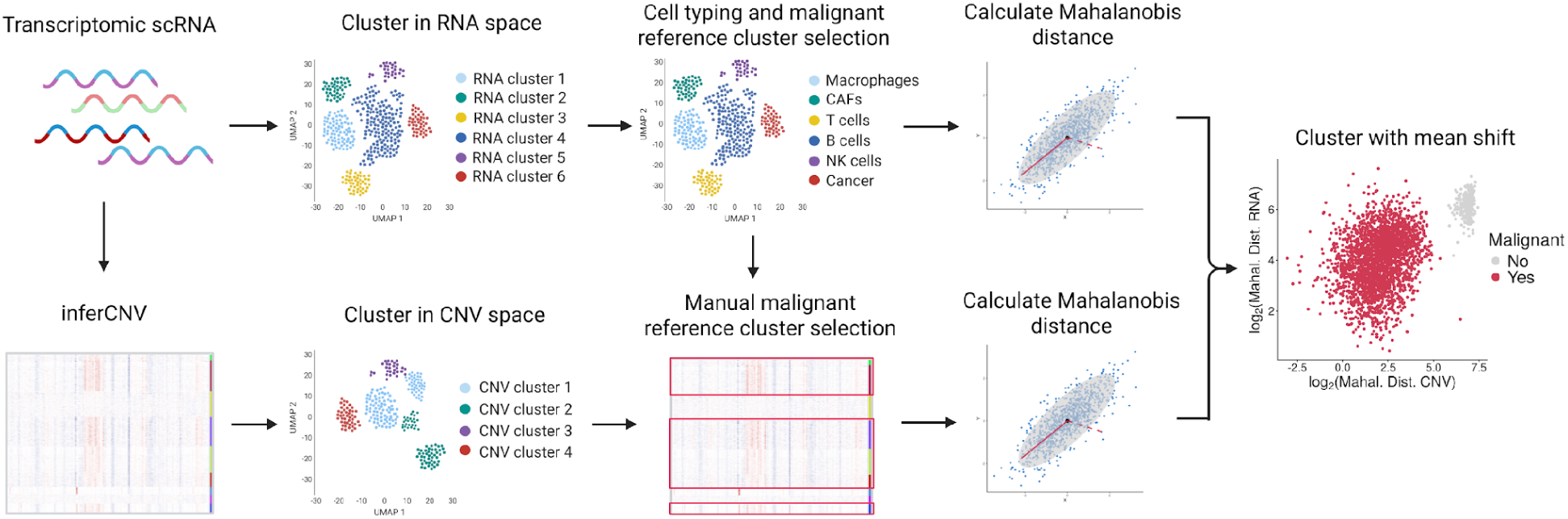
Workflow. CATFISHR malignant cell selection workflow. Two parallel procedures were developed using RNA expression and inferred CNV residual values. RNA workflow: The standard Seurat (19) SCTransform workflow was followed through clustering. Clusters and cells were then typed using SingleR (43) and scType (42). Clusters with a majority of malignant cells were selected, and the Mahalanobis distance from each cell to each reference cluster centroid was calculated. CNV workflow: Transcriptomic scRNA-seq count data were analyzed with InferCNV v1.20.0. InferCNV residual values were then used for clustering. Clusters with a majority of malignant cell types and CNV signal were selected, and Mahalanobis distances were calculated as above. For each cell, the shortest cell-to-centroid distance from each feature type was plotted, and the mean shift algorithm was used to cluster cells in this joint space. Clusters with the lowest Mahalanobis distances on both the RNA and CNV axes were selected as malignant clusters. Created in BioRender (2026) https://BioRender.com/8uh3kcq.

This approach effectively distinguishes cells with similarity in both RNA and CNV profiles from those with similarity in only RNA or only CNV space, or neither. Because CATFISHR relies solely on scRNA-seq data, it provides a simple tool to detect cancer cells in new and existing datasets, with the capacity to identify malignant cells that may be missed by cell typing or CNV inference alone. We benchmarked this approach across multiple tumor types and ground truth modalities and observed high performance in several datasets relative to cohort specific malignancy labels.

## MATERIALS AND METHODS

### Data acquisition

All analysis, except CCLONE (6), which used Python (v3.14.3), was conducted in R (v4.4.1) on a sample-by-sample basis. All datasets were filtered for samples with >500 cells. If two or more samples were present from one patient, samples with lower cell numbers were removed.

Number of samples indicated by n. Data from Pozniak (2024; n = 24) used in this paper were obtained from the KU Leuven Research Data Repository (KU Leuven RDR). The file Entire_TME.rds was used for analysis, with malignancy annotations taken from Malignant_cells.rds. Data from the Tumor Profiler scRNA-seq dataset (n = 23) were generated as detailed in Irmisch et al. and Levesque et al. (20, 21). Chronic lymphocytic leukemia (CLL; n = 3; GSE294045; (23)) and epithelial cancer datasets—specifically pancreas (GSE156405; n = 5; (24)), prostate (GSE137829; n = 5; (25)), and thyroid (GSE191288; n = 2; (26))—were obtained from sctml.xglab.tech/Download.html. One prostate sample was excluded from InferCNV CATFISHR analysis after failing CNV reference thresholds. The sample was retained in CopyKAT naive and CopyKAT CATFISHR runs. Uveal melanoma data (n = 15) were obtained from Hermes et al., manuscript in preparation. Visium data (n = 1) were generated in-house. Quality control was performed by Myriam Hofer.

### Benchmarks and Ground Truths

Benchmark or ground truth selection for each cohort (summarized in Table 1) depended on available malignant assignments. Uncertain calls, where present, were removed before analysis. Pozniak - Sophisticated Benchmark: Malignancy labels were taken from the original authors, as annotated and described in Pozniak et al. (16).

**Table 1.** Validation cohorts, benchmarks, and ground truths. Cells gives the total number of cells per cohort passing quality control; Cells analysed gives the number with a resolved benchmark or ground truth label, excluding cells with uncertain calls.

| Dataset | Tumor type | Modality | Label source | Label derivation | Cells | Cells analysed | Samples |
| --- | --- | --- | --- | --- | --- | --- | --- |
| **Ground truth** |  |  |  |  |  |  |  |
| GoT-Multi (CLL) (23) | CLL/Richter transformation | 10x scRNA-seq + targeted genotyping | SNVs/small indels | Per-cell targeted SNV mutation status | 12,327 | 3,459 | 3 |
| Tumor Profiler Visium (Lattmann et al., manuscript in preparation) | Cutaneous melanoma | 10x Visium | Histology | Selected tumor regions from histology | 4,394 | 4,394 | 1 |
| **Sophisticated benchmark** |  |  |  |  |  |  |  |
| Pozniak et al. (16) | Cutaneous melanoma | 10x scRNA-seq | Published annotations | Author-provided malignancy calls (multi-method) | 43,340 | 43,340 | 24 |
| scTML GSE156405 (Pancreas) (24) | Pancreas (PDAC) | 10x scRNA-seq | SNV-based inference | Malignancy inferred from scRNA-seq SNVs using CCLONE | 18,196 | 13,902 | 5 |
| scTML GSE137829 (Prostate) (25) | Prostate | 10x scRNA-seq | SNV-based inference | Malignancy inferred from scRNA-seq SNVs using CCLONE | 24,887 | 18,310 | 4 |
| scTML GSE191288 (Thyroid) (26) | Thyroid (PTC) | 10x scRNA-seq | SNV-based inference | Malignancy inferred from scRNA-seq SNVs using CCLONE | 6,287 | 3,878 | 2 |
| **Naive benchmark** |  |  |  |  |  |  |  |
| Tumor Profiler (TuPro) (20-22) | Melanoma (multiple subtypes) | 10x scRNA-seq | Cell typing (scROSHI) | scROSHI melanocytic cells treated as malignant | 93,927 | 93,927 | 23 |
| Uveal melanoma (UM) (Hermes et al., manuscript in preparation) | Uveal melanoma | 10x scRNA-seq | inferCNV clustering | inferCNV matrix clustering and manual selection | 74,932 | 74,308 | 15 |

**Abbreviations:** CLL, chronic lymphocytic leukemia; SNV, single-nucleotide variant; PDAC, pancreatic ductal adenocarcinoma; PTC, papillary thyroid carcinoma; scRNA-seq, single-cell RNA sequencing.

Tumor Profiler - Naive Benchmark (20–22): cells typed as melanocytes by scROSHI (v1.0.0.0; (27)) were considered malignant.

CLL - Ground Truth: CLL-RT samples were analyzed using GoT-Multi (23), which couples probe-based scRNA-seq with targeted mutation probes in the same cells. Expression-based cell typing was combined with mutation occurrence to classify cells as malignant. Eight SNVs/small indels had been analyzed by the original publication, SRRM2_T2409K, PLCG2_D993H, PLCG2_L845F, PLCG2_S1079R, B2M_E89Ter, POU2F2_K258E, JAK1_C424Ter, PRKDC_A3834E. B cells with one or more mutants detected were considered malignant, and wild-type T cells were considered normal. B cells with only wild-type calls, B cells with no mutant calls, and T cells with mutations were considered uncertain and excluded from analysis.

Pancreatic, Prostate, and Thyroid cancers - Sophisticated Benchmark: Malignancy labels were taken from CCLONE (v1.0; Python v3.14.3; (6)), using cellSNP-lite per-cell variant allele count data obtained from Li et al. (17). CCLONE infers mutation-based clonal structure and selected the optimal solution per sample by orthogonality score. Clones enriched for epithelial cells were labeled malignant and clones enriched for T/NK cells were labeled non-malignant; each cell was then assigned the label of its top-scoring clone. One prostate sample with insufficient observed SNPs (<50) was retained in analysis.

Uveal melanoma - Naive Benchmark: InferCNV was run and the residual matrix (run.final.InferCNV_obj@expr) was processed with the Seurat (v5.4.0; (19)) pipeline of RunPCA(), FindNeighbors(), and FindClusters(). Clusters with visible CNVs in heatmaps of the residual matrix were selected as malignant.

Visium - Ground Truth: samples were loaded in the 10x Loupe browser (v6.41) and spots annotated conservatively by a trained histologist (Dr. med. Maria Mannino) as Malignant or Non-Malignant based on the presence of visible tumor cells. A spot was annotated Malignant if one or more morphologically identifiable tumor cells were present within it.

### scRNA-seq data pre-processing

DecontX (v1.2.0; (28)) was used to remove ambient RNA from the filtered raw counts for all datasets except Pozniak et al. (16), which showed low ambient contamination. Using the Seurat package (v5.4.0; (19)), the standard workflow for data pre-processing and dimensionality reduction was followed. Briefly, these steps included quality control, SCTransform, normalization, scaling, feature selection, and principal component analysis (PCA). The clusterExperiment package (v2.24.0) was used to construct clusters using a wrapper for Seurat functions RunPCA(), FindNeighbors(), and FindClusters(), iterating through resolutions 0.1–0.5 in steps of 0.05 for the FindClusters() “resolution” parameter. Only variable genes were used in this step. Unassigned clusters, indicated as “-1”, were also output. Seurat was used to run UMAP dimensionality reduction and for visualizations. DoubletFinder (v2.0.6; (29)) was used to identify putative doublets in all samples. Doublet enrichment in the Hidden category relative to the other Category groups was assessed separately within each dataset using mixed-effects logistic regression implemented with glmer() in the lme4 package (v1.1-38). For each sample-category combination, the numbers of DoubletFinder-classified doublets and singlets were modeled as a grouped binomial outcome (cbind(Doublet, Singlet) ∼ Category + (1 | samples)), with samples modeled as a random intercept. Estimated marginal means and pairwise contrasts versus Hidden were computed using emmeans (v2.0.2), and reported as odds ratios. Holm-adjusted p-values and Bonferroni-adjusted confidence intervals were used for inference.

### Visium data pre-processing

Visium files were processed as in Lattmann et al., manuscript in preparation. Robust decomposition of cell types (RCTD; spacexr v2.2.1; (30)) was used for cell type identification. For each sample, 150 spots with the lowest melanoma proportions were selected. These low-melanoma spots were used as reference spots in InferCNV. InferCNV was run and the residual matrix was extracted as in scRNA-seq data. SpatialExperiment (v1.14.0; (31)) objects were used instead of Seurat. Spatially aware clustering of RNA and CNV space was performed using BANKSY (v1.0.0; (32)), with a mixing parameter λ = 0.2. Majority RCTD proportions were determined by spot. The top two clusters with highest median melanoma proportions were selected as reference clusters. CNV reference clusters were manually selected using RCTD and visual InferCNV residual expression. Mahalanobis distance calculation, mean shift clustering, and malignant cluster selection were conducted as in scRNA-seq.

### CNV Calling

InferCNV (v1.20.0) was run individually for all samples. Cells identified as T cells by both typing algorithms across all samples were used as a merged reference within a cohort. These reference T cells were removed from the query cohort. The expression matrix of InferCNV output “run.final.InferCNV_obj” was used to create Seurat objects. SCTransform was not run on these objects. Variable genes were defined as all genes with any CNV in the Hidden Markov Model (HMM) output (20_HMM_pred.repr.*.InferCNV_obj). These Seurat objects were clustered using clusterExperiment and the procedure described above with resolutions in the range 0.25–0.70 in increments of 0.05.

CopyKAT (v1.1.0) was run on pancreatic, prostate, thyroid, and CLL cohorts, with results processed generally as described above, except for changes mentioned below. All cells labeled as “T/NK cells” within a sample were selected as normal references in the CNV caller. CopyKAT bin-level CNA output (\*_CopyKAT_CNA_results.txt) was assembled into a bins × cells matrix and residualized by subtracting, for each bin, the mean value across a set of normal reference cells (≥ 5). Bin intervals were approximated from adjacent chromosomal positions within each chromosome, and genes (HGNC; chrom, start_position, end_position) were assigned to overlapping bins. Gene-level residual CNV values were computed as the mean residual across bins overlapping each gene, yielding a genes × cells matrix which was used to create a Seurat object. These Seurat objects were clustered using clusterExperiment as above. The top variable genes, with an upper threshold of 500, were selected by variance of residual CNV signal for downstream analysis. In mean shift clustering the RNA space did not change. Prostate SRR10162350 was filtered out in the InferCNV cohort due to not passing the CNV cluster reference criteria but was not filtered out in CopyKAT.

### Cell Typing

Two cell typing algorithms were run on each sample. SingleR (v2.6.0; (43)) was run using the reference dataset and cell annotations from Tirosh et al. (15) for all selected samples from the cutaneous enriched melanoma datasets Pozniak (16) and Tumor Profiler (20, 21); the celldex R package (v1.14.0) HumanPrimaryCellAtlasData(legacy = TRUE) reference (37) was used for Pancreas, Prostate, Thyroid, and CLL; and Durante et al. (38), selected using TISCH (39, 40), was used as a reference for UM.

scType functions were taken from GitHub (42). Input marker lists are listed in Supplementary Data 1. Markers for cutaneous enriched melanoma datasets Pozniak and Tumor Profiler were manually constructed using the literature; Pancreas markers from Peng et al. (33); Prostate markers from Dong et al. (34); Thyroid markers from Pu et al. (35); UM markers from Lin et al. (36). An aggregated cell type for each cell was identified by matching outputs from each algorithm.

### Cluster Selection in RNA and CNV space

Clusters to be designated as cancer cell type clusters were manually selected in CNV space using heatmaps of residual expression (from InferCNV output “run.final.InferCNV_obj”), HMM InferCNV output

(“20_HMM_pred.repr_intensitiesHMMi6.leiden.hmm_mode-subclusters.Pnorm_#.InferCNV_obj”) heatmaps, and cell typing. Clusters with visible CNVs that also had a majority of cells typed as the cancer cell type by both typing algorithms were chosen as “cancer cell clusters”. One Pozniak sample (16) with CNVs only in the unassigned cluster was removed. In RNA space, clusters with a majority of cells typed by SingleR or cluster cell type in scType as the cancer cell type were selected.

### Calculating Mahalanobis Distance

Mahalanobis distance in PCA space was calculated for each cell to each cluster centroid. A small ridge penalty (λ = 1e-6) was added to the diagonal of each reference cluster covariance matrix to ensure numerical stability. The workflow below was conducted separately in RNA and CNV space. The number of PCs retained was determined by the consecutive difference in variance explained. PCs were retained up to the last PC where the difference between consecutive PCs exceeded 0.1%. Cluster centroids were calculated in PCA space by taking the median of each PC. The cluster centroid for the unassigned cluster (“-1”) was not calculated. The covariance matrix and Mahalanobis distance were calculated for each cluster using the stats package (v4.4.1).

### Mean Shift

The smallest Mahalanobis distance between a cell and any of the selected malignant clusters was used to run the mean shift algorithm (meanShiftR v0.56). Each cell therefore had a Mahalanobis distance associated with it in RNA and CNV space. Distances were log₂ transformed. Mean shift was run for each sample over bandwidths 0.4–0.8 in steps of 0.1 for 500 iterations. The maximum number of possible clusters was 7. The silhouette distance (cluster v2.1.8.1) of each run was calculated, and the model with the highest mean silhouette distance was used in further analysis. Malignant clusters were identified as the clusters with the smallest distances in both RNA and CNV space. Balanced accuracy and F1 score were calculated using yardstick (v1.3.2). Cohen’s d was calculated using the rstatix package (v0.7.3).

### Cancer Metaprograms

Within each dataset, 46 cancer metaprogram signatures from the Weizmann Institute (4, 41) were scored using UCell (44) and scaled (Supplementary Data 2). Category-associated variability was quantified using the Kruskal–Wallis ε² effect size, together with the interquartile range, to exclude non-differentiating signatures. For each signature, median scores were computed per category. Euclidean distances were then calculated between the Hidden or Missed median and the Malignant and Non-Malignant medians to quantify overall similarity in signature space.

## RESULTS

### Malignant cell selection by single modalities

The Tumor Profiler (TuPro) scRNA-seq dataset (n=23 samples; n=93,927 cells; (20, 21)) is a dataset consisting of several melanoma subtypes. The published cell type annotations (scROSHI (27)) of TuPro were evaluated using two additional transcriptome-based cell typing algorithms: SingleR (43), which provides per-cell annotations, and scType (42), which provides cluster level annotations. Discordance in cell type calls across the three annotation sources was summarized in Figure 2A. Coarse cell type labels for 62.7% of cells agreed across all algorithms (high-confidence calls), and 3.9% of cells received a different label from each method.

**Figure 2.**
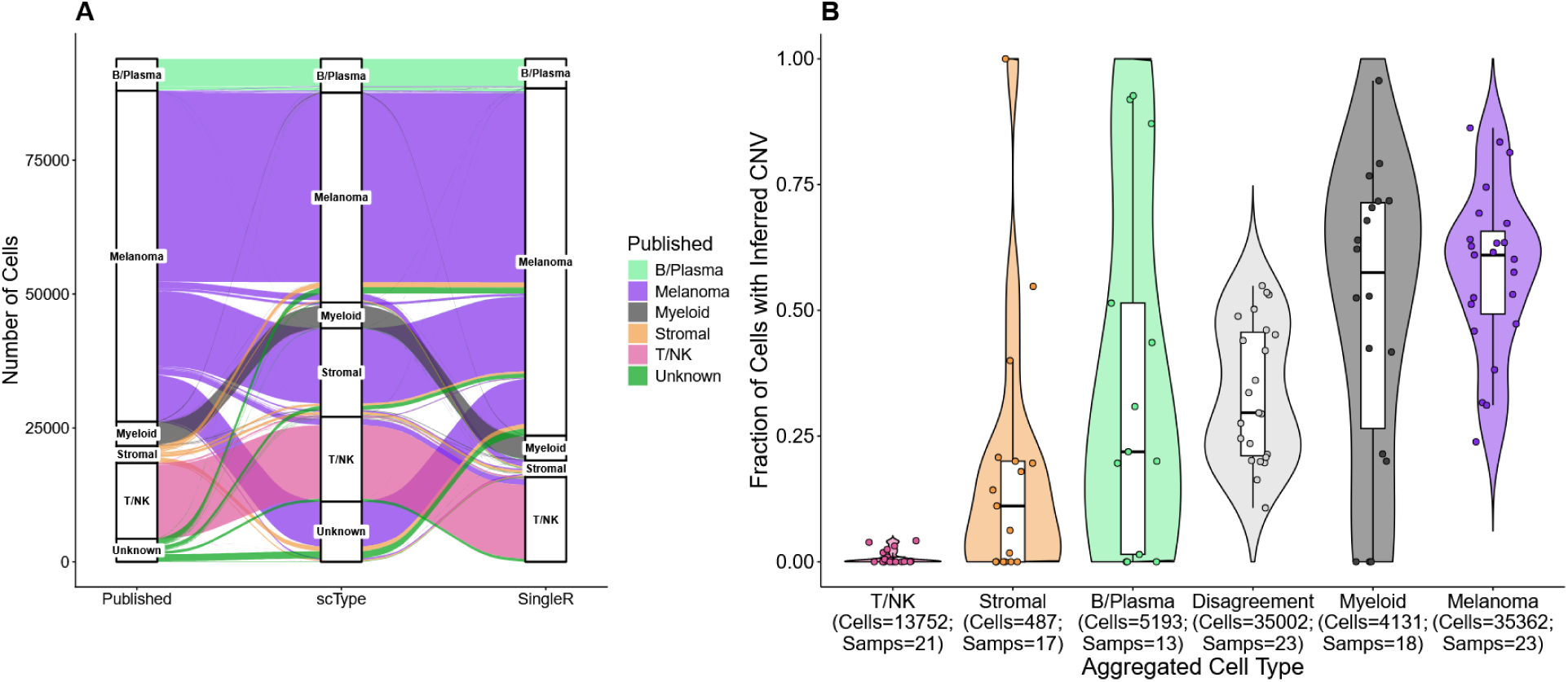
Coarse cell typing and inferred CNV comparisons. A) Alluvial plot showing Tumor Profiler cells (n = 93,927) across all samples (n = 23). Strata indicate coarse cell typing by SingleR (43), scType (42), and published annotations (scROSHI, (27)) obtained from the Tumor Profiler study authors. Alluvia are colored by the Published cell type. B) Cell CNVs were labeled at the chromosome level. The fraction of CNV positive cells among high-confidence cell types (same coarse label across scType, SingleR, and published annotations) is shown, with each point indicating a sample. Not all samples have every cell type. Numbers of cells and samples in each cell type are indicated in the axis names. Cells with any disagreement across the three annotation sources are labeled “Disagreement”.

Published immune-cell labels showed high agreement across all methods for T/NK (97.2%), myeloid (91.0%), and B/plasma cells (87.3%). However, high confidence consensus was lower for melanoma cells (57.2%) and stromal cells (15.2%). Inferred CNV signals from InferCNV, summarized by chromosome as the presence or absence of at least one gene with a high-confidence amplification or deletion, was used to label cells as CNV positive (any CNV detected) or CNV negative (no CNVs detected across all chromosomes) and compared to high confidence cell type calls. (Figure 2B). Among cells typed as non-melanoma across all three cell typing algorithms, 17.7% were CNV positive, while 44.8% of those typed as melanoma were CNV negative. These observations motivated the integration of each modality in CATFISHR to increase confidence in malignant cell selection.

### CATFISHR reference cluster selection

Malignant reference clusters in RNA and CNV space were selected conservatively, to ensure high confidence malignant references in downstream analyses, while all remaining cells were treated as query cells of an initially undefined malignant status (Figure 3). Cluster selection was performed on a sample-by-sample basis. CNV clusters were generated on the InferCNV residual expression matrix and plotted as a heatmap which was then used to select reference clusters. Selected clusters (dark blue in Cluster Type, Figure 3A) required visible, consistent CNV patterns within the cluster and majority agreement across cell typing methods for the tumor cell-of-origin type. Both cell typing and CNV validation is required as CNV space manually selected reference clusters were additionally filtered to ensure clusters contained a majority of the cancer cell type according to both scType and SingleR. Selection of RNA space reference clusters required the same majority cell type criterion (Figure 3B). A UMAP representation of RNA space query and reference clusters shows the relationship between the clusters (Figure 3C).

**Figure 3.**
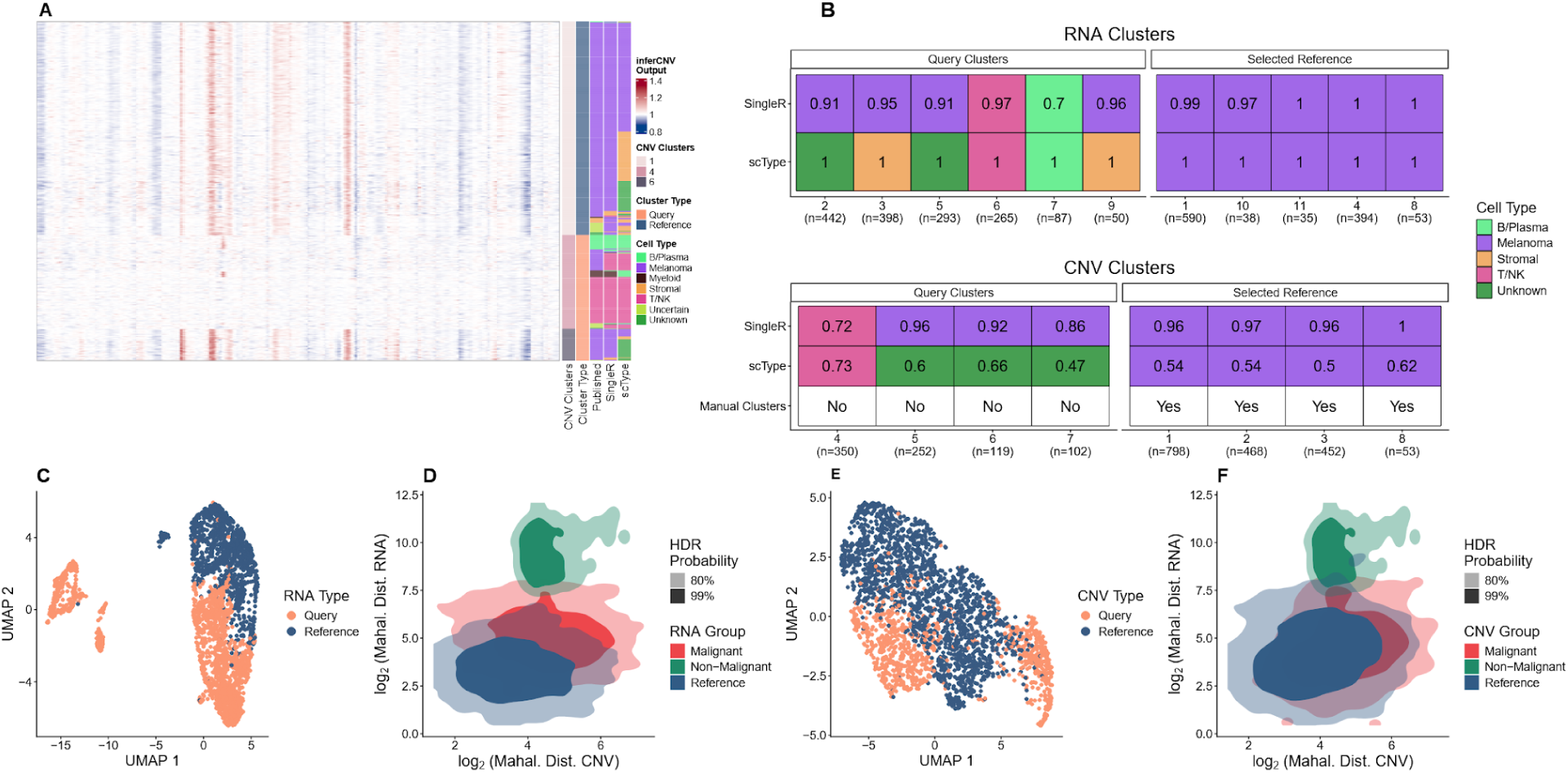
Reference Cluster Selection. Analysis of an example Tumor Profiler sample is shown. A) InferCNV residual expression heatmap subset. Row annotations show CNV space clusters (“CNV Clusters”), cluster designation (“Cluster Type”; Reference vs Query), and coarse cell type labels from three annotation sources. “Reference” indicates clusters selected based on clear CNV patterns and concordant cancer cell-of-origin typing across methods (high confidence malignant); all other clusters are treated as “Query”. B) RNA and CNV space reference cluster selection and filtering based on majority cell typing. For each cluster, tile values indicate the proportion of the majority cell type, and tile colors indicate the majority cell type. Clusters are grouped by whether they were ultimately selected as reference clusters. In CNV space, “Manual Clusters = Yes” indicates clusters visually selected from InferCNV heatmaps prior to majority typing filtering. Cell counts per cluster are shown on the x-axis. C) RNA UMAP showing reference clusters and query cells. D) CATFISHR output in RNA–CNV Mahalanobis distance space. Axes show log₂ distances to the nearest RNA and CNV reference clusters. RNA reference cells are shown in blue; query cells classified as “Malignant” are shown in red and “Non-Malignant” cells in grey. Shaded regions indicate highest density regions (HDR; 80% and 99%). E–F) Corresponding CNV space UMAP (E) and Mahalanobis distance plot with CNV reference clusters (F) for the same cells.

The mean shift algorithm is used to separate malignant and non-malignant compartments, after calculation of the shortest Mahalanobis distance between reference cluster centroids to all cells in RNA and CNV space. Maximum mean silhouette distances were used to select the optimal mean shift bandwidth, which varies by cluster numbers (Supplementary Figure 1). Mean shift output using the selected RNA reference clusters is shown in Figure 3D, including the resulting malignant and non-malignant calls in RNA space. The shortest Mahalanobis RNA reference centroid-to-cell distance is plotted on the y-axis. Corresponding plots in CNV space for the same sample and cells are shown in Figure 3E–F, highlighting the CNV reference clusters. The shortest Mahalanobis centroid-to-cell distances for CNV reference clusters are plotted on the x-axis.

### CATFISHR output

The CATFISHR output displaying the orientation of malignant and non-malignant clusters for the full TuPro cohort (n=23 samples; n=93,927 cells) is shown in Figure 4A. Cell density is summarized using high-density regions (HDR). Cell clusters with low distances in both RNA and CNV space were classified as “Malignant” (red) comprising 70.2% of cells (n = 65,904). Clusters with high distances in both spaces were “Non-Malignant” (green) comprising 29.8% of cells (n = 28,023). Mean shift malignancy calls (red cluster in Figure 4A) were compared to the cohort benchmark derived from published scROSHI annotations (27) (Figure 4B). Cells typed as “Melanoma Melanocytic” or “Melanoma Mesenchymal” were considered malignant. The overlap categories “Malignant” (63.2%) and “Non-Malignant” (27.3%) indicate agreement between CATFISHR and benchmark calls. “Missed” (2.5%) cells were labeled malignant by the benchmark but not by CATFISHR, whereas “Hidden” (6.9%) cells were labeled malignant by CATFISHR but not by the benchmark. Hidden cells were shown to have visible CNVs, while Missed cells showed very few to no CNVs (Supplementary Figure 2).

**Figure 4.**
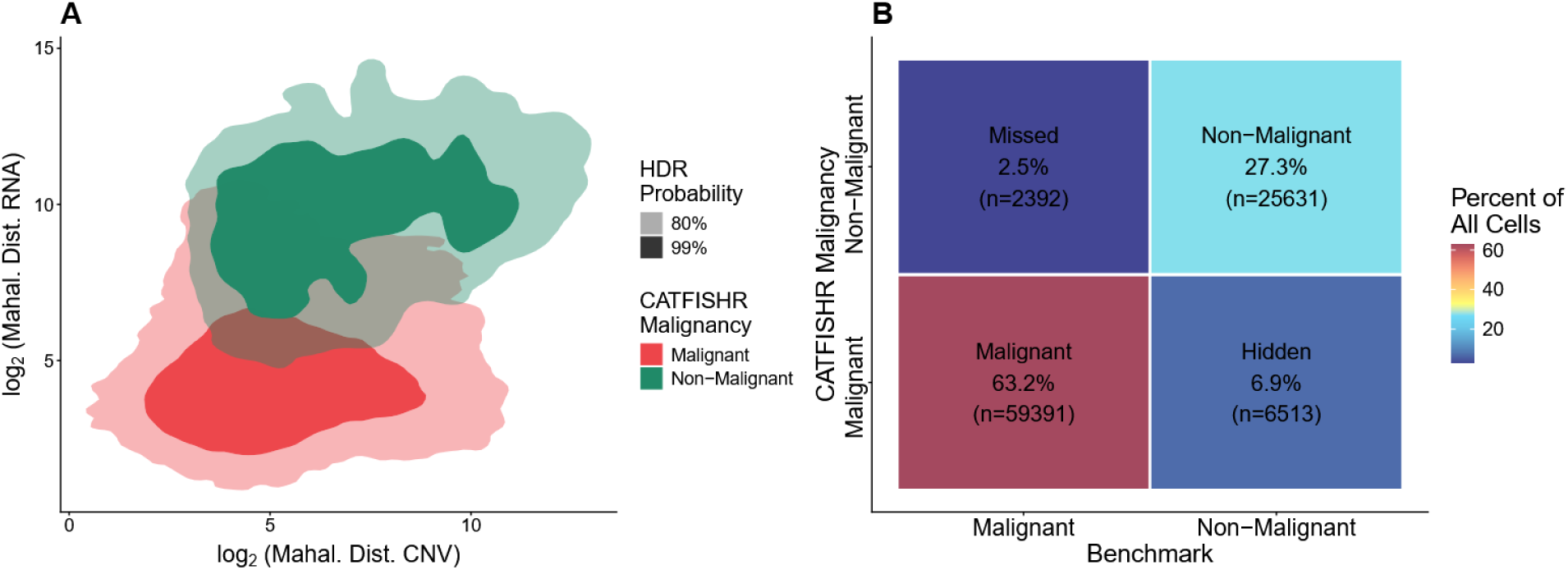
Tumor Profiler CATFISHR Output. A) RNA and CNV Mahalanobis distance space for the full Tumor Profiler cohort (n=23 samples; n=93,927 cells) showing CATFISHR malignancy calls. Axes show log₂ Mahalanobis distances in RNA and CNV space (using the minimum distance to the selected reference clusters). Cell density is shown by highest density regions (HDR; 80% and 99%). B) Agreement between CATFISHR malignancy calls and the benchmark. benchmark was defined using scROSHI cell typing (27): cells typed as melanocytic were considered malignant. Tiles are colored by the percentage of all cells in each compartment and annotated with the percentage and cell count. “Malignant” and “Non-Malignant” indicate agreement. “Missed” indicates benchmark malignant cells called non-malignant by CATFISHR. “Hidden” indicates cells called malignant by CATFISHR but Non-Malignant by the benchmark.

### Validation in other scRNA-seq datasets

To evaluate CATFISHR across cancer types and benchmark/ground truth modalities, we analyzed multiple datasets with available malignancy labels (Table 1). Full details of benchmark/ground truth definition for each cohort are provided in the Methods section.

Performance was assessed per sample using balanced accuracy and F1 score, stratified by benchmark/ground truth source (Figure 5A). Published annotations were used as provided by the original studies without modification unless noted otherwise (Table 1; Materials and Methods). Cells without published annotations were removed from analysis. The median balanced accuracy was above 0.85 for 5/7 scRNA-seq datasets, and median F1 score above 0.80 for 5/7 scRNA-seq datasets. Across all scRNA-seq cohorts (76 samples, 251,124 cells), of cells which disagreed between benchmarks and CATFISHR malignancy calls, 8.3% were classified as Hidden and 1.3% as Missed. Cells which agreed on malignancy calls between benchmark and CATFISHR were 50.6% Malignant and 39.8% Non-Malignant. However, the proportion of categories varied across the datasets (Supplementary Figure 3). The highest confidence orthogonal ground truth available was for the CLL cohort via targeted mutation probes (Figure 5B). CATFISHR and SNV/small indel based ground truth calls showed high agreement, with 5.3% of all cells Missed and 5.3% Hidden. Across all CLL cells, this corresponds to median balanced accuracy of 0.89 and median F1 of 0.89.

**Figure 5.**
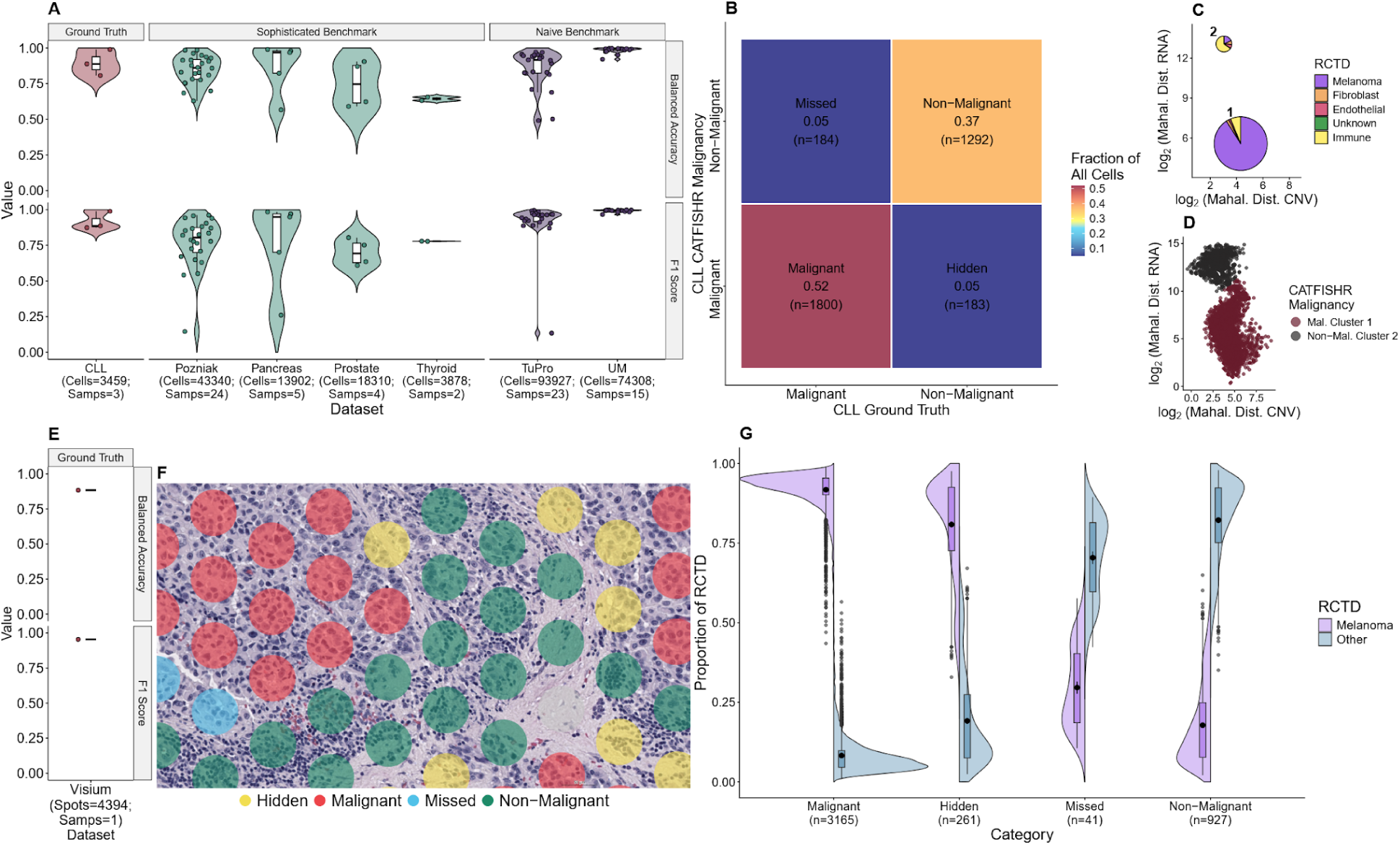
Benchmark/Ground Truth Comparisons. A) Concordance of CATFISHR malignancy calls with available benchmarks/ground truths (benchmark/ground truth confidence indicated by the top facet). On the horizontal facets, balanced accuracy (top) and F1 score (bottom) were computed per sample; points represent samples and violins show the distribution within each dataset. The number of cells and samples per dataset is shown on the x-axis labels. B) Benchmark/ground truth comparison matrix for the CLL cohort, which uses an orthogonal SNV/small indel based ground truth, the highest confidence ground truth in this study. Labels as in Figure 4B. C–D) Example CATFISHR output for 10x Visium data. C) RCTD estimates summarized per mean shift cluster; pie radii are scaled by the number of spots contributing to each cluster (cluster 1: n = 3426, 2: n = 968). D) CATFISHR malignancy calls for the same spots and clusters. E) Balanced accuracy and F1 score for the Visium sample. F) Subsection of the Visium sample showing spot level agreement between CATFISHR and histology based ground truth. G) RCTD deconvolution proportions of melanoma versus all other cell types stratified by overlap category for the Visium sample (n = 1), with 4394 spots.

To assess whether Hidden calls could be explained by technical artifacts, we compared standard scRNA-seq QC metrics across overlap categories (Supplementary Figure 4). For cohorts with more than 3 samples, QC differences between Hidden and each other category were summarized using Cohen’s d computed on per-sample median QC values. Results showed inconsistent association of QC values with any category (Supplementary Table 1). Additionally, the proportion of cells identified as doublets for each category varied across the datasets (Supplementary Figure 5). To identify doublet-enrichment in Hidden groups, mixed-effects logistic regression was run for each dataset. Estimated marginal means and pairwise contrasts versus Hidden were then computed and reported as an odds ratio (Supplementary Table 2). Datasets with fewer than 3 samples were excluded from analysis. Doublet enrichment varied by cohort, with all Hidden being enriched for doublets from all comparisons in Pancreas, and Malignant vs Hidden in Pozniak. All other comparisons in all datasets were not significant or the Hidden category was not enriched for doublets.

Ambient RNA enrichment was additionally assessed by comparing median DecontX contamination in Hidden cells against other categories at the project level. Hidden cells did not show consistent enrichment relative to Malignant or Non-Malignant cells, with differences varying across datasets and generally remaining small. In contrast, Hidden cells showed lower ambient RNA enrichment than Missed cells across all datasets, indicating that Missed cells had the highest ambient RNA contamination among the evaluated categories. (Supplementary Figure 6). Expression profiles of categories were tested by calculating cancer metaprogram signatures for each cell using Weizmann Institute signatures (4, 41) and UCell (44). Category medians were calculated in signature space and the distances compared in Supplementary Figure 7. Comparisons of Hidden and Missed versus Malignant and Non-Malignant categories showed the Malignant category closer to the Hidden cells than the Non-Malignant category in 5/7 cohorts. On the other hand, Missed cells were closer to Non-Malignant than Malignant in 5/7 of cohorts.

### Validation in Visium data

One sample from the TuPro project was spatially analyzed using 10X Visium, by treating spots analogously to cells and applying the above workflow adapted for spatial data (Materials and Methods). Robust Cell Type Decomposition (RCTD) (30) was used analogously to cell typing during reference selection. The CATFISHR output for this sample, aggregated by cluster, is shown in Figure 5C. The unaggregated version of the sample with individual spots is shown in Supplementary Figure 8A. Two mean shift clusters were identified and summarized by their RCTD cell type composition, showing cluster 1 as highly melanoma enriched and cluster 2 as highly immune enriched. These match the corresponding CATFISHR malignancy calls as shown in Figure 5D (Supplementary Figure 8B).

Visium balanced accuracy was 0.88, and F1 score was 0.95 (Figure 5E). In a representative Visium region, spot level overlap categories (“Malignant”, “Non-Malignant”, “Hidden”, “Missed”) are shown in Figure 5F. RCTD cell composition pie charts for each spot and the associated category label are shown in Supplementary Figure 9. Missed spots were rare (0.9% of all spots), while hidden spots comprised 5.9% of all spots (Figure 5G). Hidden spots showed a small increase in RCTD melanoma proportion compared with Missed spots. Melanoma proportions were slightly higher in Hidden versus Missed spots (median 0.87 vs median 0.28), with a corresponding marginal decrease in “Other” (median 0.13 vs 0.72). For the sample, histology-based spot annotation was conservative, resulting in relatively few Missed spots (n = 41) and a larger Hidden compartment (n = 261).

### Comparison with CopyKAT Native and CopyKAT input into CATFISHR

To assess robustness across CNV callers, we repeated the analysis using CopyKAT CNV output as input to the CNV space component of the CATFISHR workflow. CopyKAT required sample specific reference cell selection and preprocessing to generate a gene-level residual CNV matrix; downstream steps were otherwise identical to the InferCNV input CATFISHR workflow (Materials and Methods). We evaluated CopyKAT input CATFISHR malignancy calls and native CopyKAT malignancy calls against cohort ground truths in the CLL, pancreas, prostate, and thyroid datasets using balanced accuracy and F1 score per sample (Figure 6). For comparison, InferCNV input CATFISHR performance from Figure 5A is replotted. The median differences in balanced accuracy and F1 score for each cohort are summarized in Supplementary Table 3. The difference between the CATFISHR runs was small for several samples, particularly in Pancreas and CLL, with balanced accuracy considered equivalent in 5/14 comparison-evaluable samples and F1 score considered equivalent in 2/14 comparison-evaluable samples. Results were more consistent in Pancreas and CLL, whereas Prostate and Thyroid showed larger shifts in at least one comparison. Neither CATFISHR algorithm consistently outperformed native CopyKAT: using aggregated comparison-specific sample counts, either CATFISHR run performed better than native CopyKAT in 7/29 balanced accuracy comparisons and 9/29 F1 score comparisons. Overall, median differences were generally modest, indicating the algorithms performed similarly.

**Figure 6.**
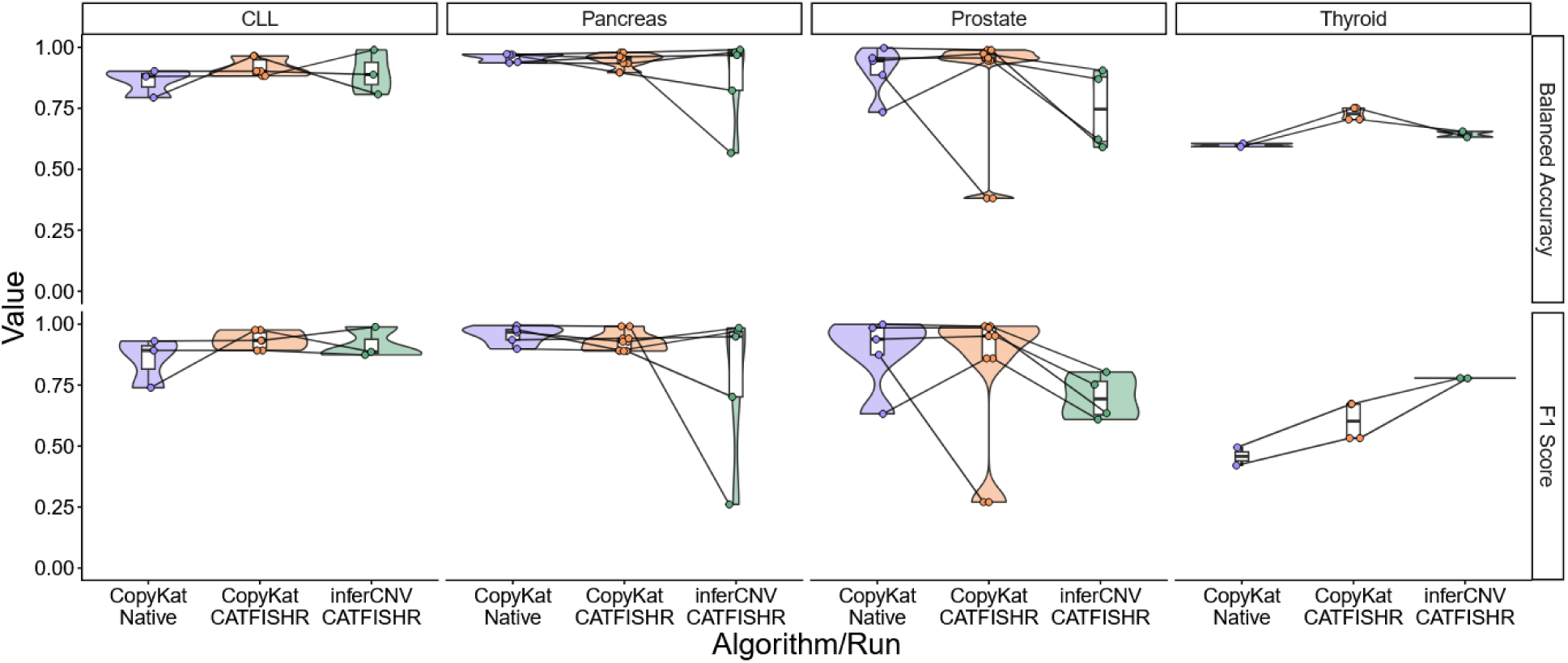
CopyKAT matrix as Input to CATFISHR. CopyKAT was run per sample, with T cells used as normal/reference cells. Preprocessing of CopyKAT matrix described in Materials and Methods. Cohort benchmark/ground truth compared across three approaches: native CopyKAT calls, CopyKAT input CATFISHR calls, and InferCNV input CATFISHR calls. ‘Uncertain’ calls were excluded. Lines connect the same sample across algorithms/runs. One sample in the Prostate cohort was excluded from the InferCNV input CATFISHR results due to failing CNV reference cluster criteria, but was retained for both CopyKAT analysis.

## DISCUSSION

Across diverse datasets and benchmarks, CATFISHR showed strong performance in several cohorts, including the CLL dataset with orthogonal SNV-based labels, which provides the highest-confidence scRNA-seq evaluation in this study. Interpretation of these metrics depends on the nature of the available malignancy labels; both F1 score and balanced accuracy treat Hidden cells as false positives against the benchmark, meaning reported scores represent a conservative lower bound on true performance. Despite this, high metric values in most datasets, low proportions of Missed cells, and the similarity of Hidden cells to Malignant in signature space collectively indicate that CATFISHR captures malignant cells more comprehensively than the benchmarks alone. Tested datasets span orthogonal SNV-based genotyping (CLL), histology-defined tumour regions for spatial data (Visium), and multiple forms of published annotations derived from RNA, inferred CNV, and inferred SNV-based evidence.

Annotations were therefore separated by confidence, and non-orthogonal sources were treated as benchmarks rather than definitive truth. In particular, cohorts in which malignancy labels were derived from CNV-based pipelines are informative for agreement and practical utility but do not provide fully independent validation of CNV-driven methods. In contrast, the CLL cohort provides a stringent evaluation because malignancy is defined by targeted detection of somatic mutations in the same cells. CATFISHR achieved a median balanced accuracy of 0.89 and F1 score of 0.89 in this cohort, with Hidden and Missed cells each comprising 5.3% of all cells, supporting the reliability of the framework’s malignancy calls when evaluated against independent ground truth.

Cells/spots uniquely identified as malignant by CATFISHR (“Hidden”) were observed across datasets. Consistent CNV patterns in CNV space visualizations supported the interpretation that a subset of Hidden cells represent malignant cells, whereas Missed cells showed little to no CNV signal. Moreover, comparisons in cancer metaprogram signature space indicated Hidden cells in most datasets had profiles closer to Malignant than Non-Malignant profiles. Hidden cells likely represent a heterogeneous group, including malignant cells missed by benchmark annotations, cells with borderline malignant transcriptional/CNV profiles, and potentially a smaller fraction of technical or annotation-related discordances. Their similarity to Malignant cells in CNV and metaprogram space supports a biological basis for many Hidden calls, but orthogonal validation is needed to distinguish these possibilities definitively.

QC comparisons across overlap categories, including effect-size analysis in cohorts with sufficient sample numbers, did not reveal systematic QC differences that could explain Hidden calls. Hidden cells were also not consistently enriched for doublets or ambient RNA compared to other categories across all datasets. Missed cells however were enriched for ambient RNA compared to Hidden for all cohorts. The enrichment of ambient RNA among Missed cells, together with weaker CNV signal, suggests that some Missed calls may reflect cells with degraded or contaminated transcriptional profiles, low malignant signal, or ambiguous RNA/CNV evidence rather than systematic failure of the framework alone. Interpretation of the Thyroid cohort should be cautious as it had the highest average estimated ambient RNA contamination across cohorts (data not shown), in addition to limited sample numbers.

In Visium data, Hidden spots showed modestly higher melanoma proportions than Missed spots by RCTD deconvolutions. Because Visium spots capture mixtures of cell types and tumor cells can be difficult to identify visually at spot resolution, discordance between histology-based labels and transcriptional deconvolution is expected in some cases. These results suggest that a subset of Hidden spots may contain melanoma signals that were not called by histology. Although characterization is beyond the scope of this study, the presence of Hidden cells across datasets raises the possibility that standard annotation pipelines systematically exclude a subset of malignant cells, warranting further investigation.

We also tested robustness to CNV caller choice by substituting CopyKAT-derived matrices into the workflow. Performance varied by sample and cohort, and neither CNV input source consistently outperformed the other. This suggests that CATFISHR’s primary contribution is the integration of RNA and CNV information rather than improvement over any individual CNV caller. Nevertheless, compatibility with alternative CNV inputs expands the framework’s flexibility and supports its use across datasets where CNV caller performance may differ.

Several limitations should be noted. CATFISHR depends on coherent structure in both RNA and CNV space, and performance may be reduced in cancer types with generally low-to-moderate CNV burden: pancreatic ductal adenocarcinoma (PDAC), prostate adenocarcinoma, and papillary thyroid carcinoma (PTC) (45). Additionally, individual sample performance varied and that extreme cases were driven by different factors including class imbalance, sample quality, and sample-specific biology. Moreover, because CNV profiles are inferred from transcript abundance, RNA-based CNV approaches may be less sensitive in cells with low transcriptional activity, low RNA content, or weak copy-number signal. As a result, quiescent or transcriptionally subdued malignant phenotypes may be under-detected by CATFISHR and related approaches. Conversely, non-malignant myeloid cells showed inferred CNV frequencies comparable to melanoma cells, consistent with lineage-specific expression programs and immune-cell genomic rearrangements such as V(D)J recombination complicating inferred CNV output (14).

In addition to these technical constraints, manual selection of CNV reference clusters and mean shift clusters introduces subjectivity, and some cohorts have limited sample numbers and/or lower confidence benchmarks, which complicates interpretation. Future work could focus on automating reference selection, improving uncertainty reporting (e.g., an explicit ‘uncertain’ category) such as with silhouette scores, or Hierarchical Density-Based Spatial Clustering of Applications with Noise (HDBSCAN) (46), and incorporating additional orthogonal signals where available. Despite these limitations, CATFISHR performed comparably across CNV input sources, achieved high F1 and balanced accuracy scores in most datasets, and identified malignant-like spots in spatial data that were not captured by histology annotation. Hidden cells frequently resembled Malignant cells in profile space and did not show systematic QC differences or consistent enrichment for doublets or ambient RNA. Conversely, Missed cells comprised less than 5.4% of cells in all projects, were enriched for ambient RNA relative to Hidden cells, and more closely resembled Non-Malignant profiles in many datasets. Together, these findings support CATFISHR as a flexible framework for malignant cell identification from scRNA-seq data, particularly when benchmark annotations are incomplete or rely partly on non-orthogonal evidence.

## ACKNOWLEDGEMENTS

The investigators are grateful to all patients and their families whose donated material made this study possible. They also thank all contributors who aided in the generation, analysis, interpretation, and development of ideas, in particular Irene Keller, Prof. Dr. med. Michael Krauthammer and Prof. Dr. Valentina Boeva.

## AUTHOR CONTRIBUTIONS

Prachi Shukla: Conceptualization, Data Curation, Formal analysis, Investigation, Methodology, Software, Validation, Visualization, Writing—original draft, Writing—review & editing. Maria Mannino: Investigation, Myriam Hofer: Data Curation, Evelyn Lattmann: Data Curation. Veronika Haunerdinger: Supervision, Writing—review & editing. Patrick Turko: Conceptualization, Methodology, Supervision, and Writing—review & editing. Mitchell P. Levesque: Supervision.

## SUPPLEMENTARY MATERIAL

Supplementary Materials are available with the online version of this preprint.

Supplementary Figures 1-9 and Supplementary Tables 1-3 are available online as Supplementary Material.

Supplementary Data 1: scType cell type marker genes (SupplementaryData1.xlsx).

Supplementary Data 2: Weizmann Institute cancer metaprograms gene signatures (SupplementaryData2.xlsx).

## DATA AVAILABILITY

A R package implementing CATFISHR Mahalanobis distance calculation and mean shift clustering steps can be found at github.com/LevesqueLabHub/CATFISHR. A frozen version corresponding to this manuscript is archived in Zenodo at https://doi.org/10.5281/zenodo.20530569. Public datasets analyzed in this study are available from the repositories and accession numbers listed in Table 1 and the Methods section. Supplementary Data 1 and Supplementary Data 2 are available with the online version of this preprint.

