## Supplementary Material for "CATFISHR: integration of transcriptome and inferred CNV profiles for comprehensive malignant cell identification"

### Supplementary Figure 1

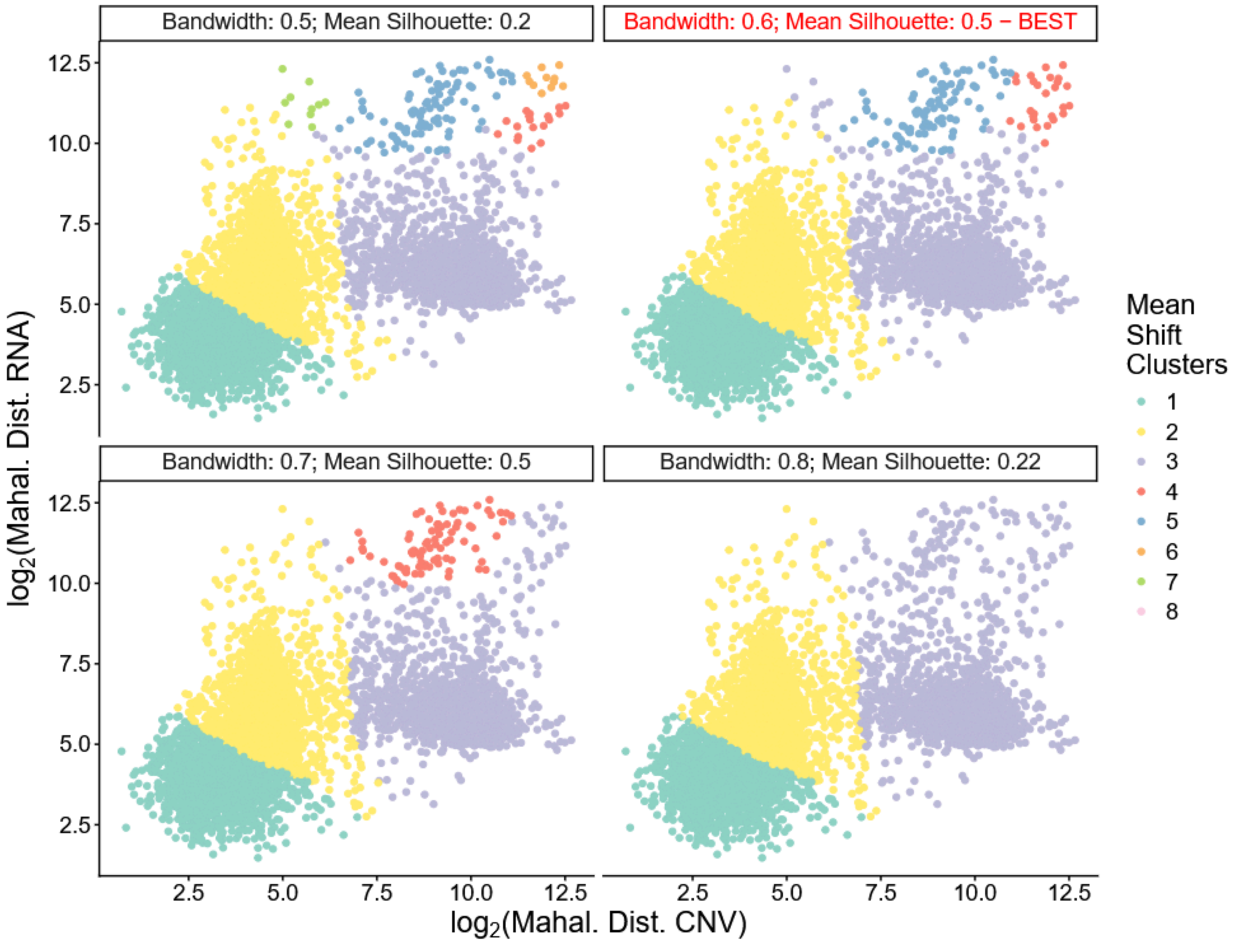


**Supplementary Figure 1.** Mean shift bandwidth selection. Example sample mean shift clustering across bandwidths 0.5-0.8. Bandwidths and mean silhouette distances are indicated in the facets. The best bandwidth is selected by the maximum mean silhouette distance, shown in the facet with the red title.

### Supplementary Figure 2

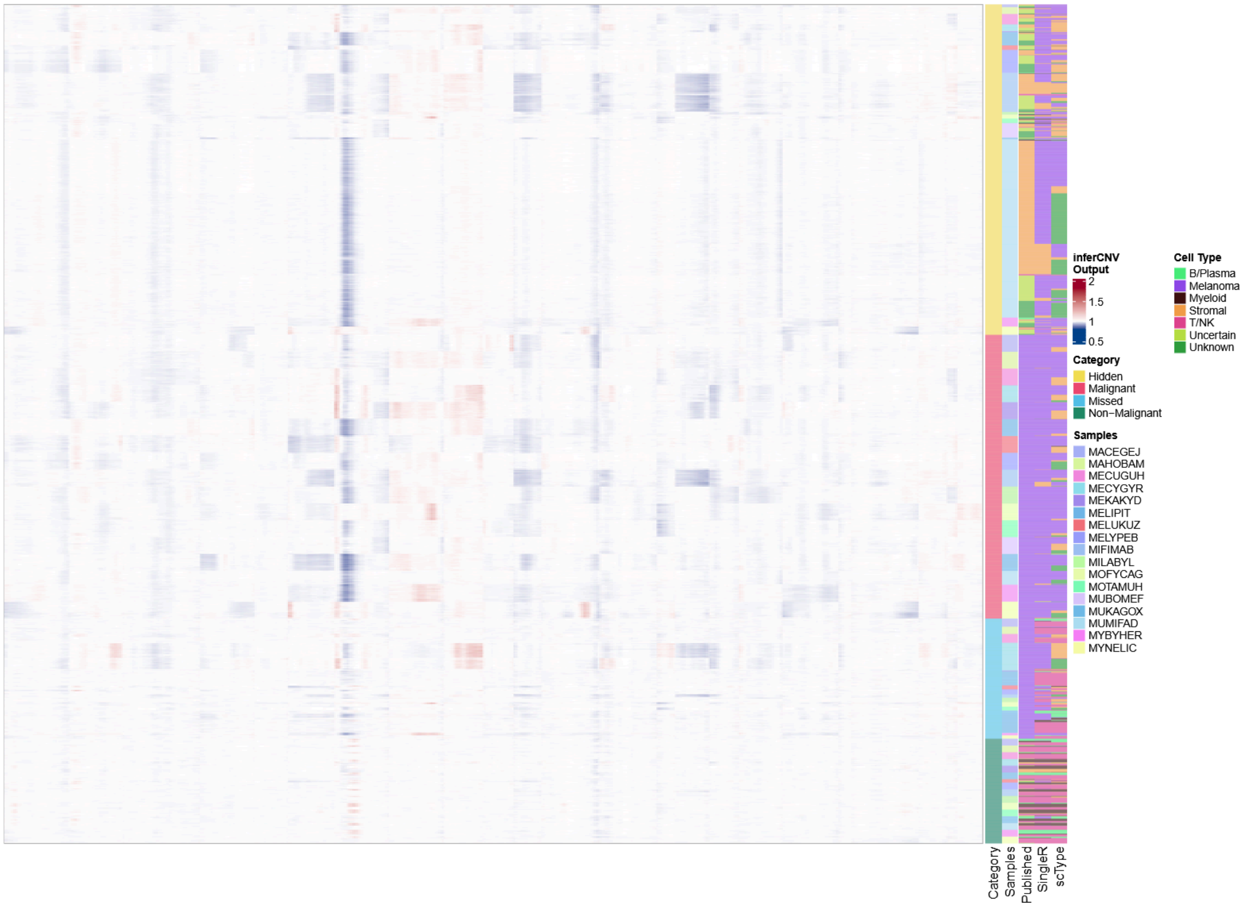


**Supplementary Figure 2.** Heatmap of Tumor Profiler Hidden cell CNV residual expression. Heatmap showing all Hidden and Missed cells from the Tumor Profiler cohort and a random selection of Malignant and Non-Malignant cells.

**Supplementary Figure 3**

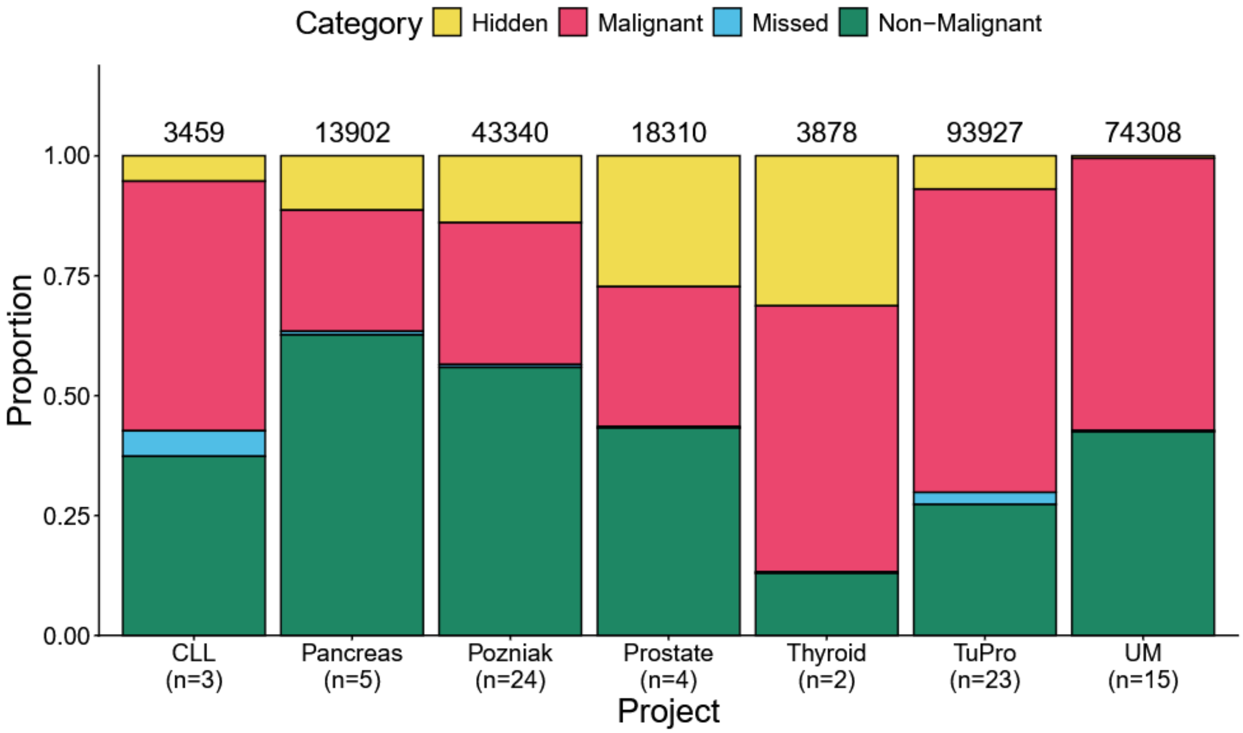


**Supplementary Figure 3.** Project overlap category compositions. Proportion of overlap categories for each project. The number above the bars indicates the total number of cells in a project. Sample numbers are shown in the x-axis.

**Supplementary Figure 4**

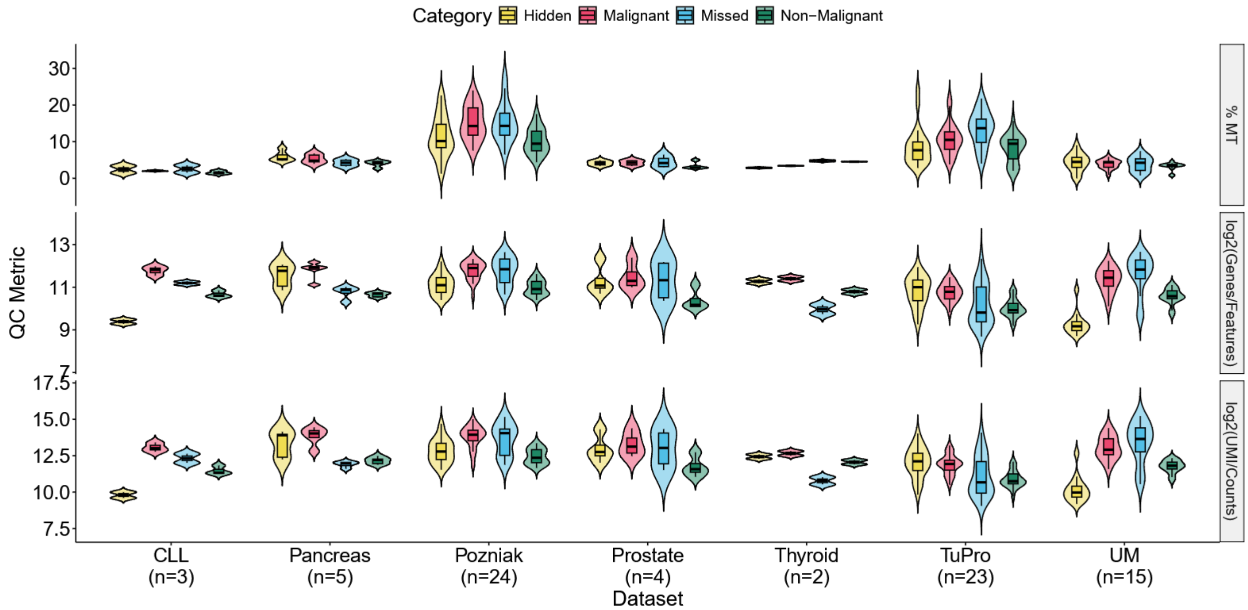


**Supplementary Figure 4.** Quality metrics in CATFISHR vs ground truth overlap categories. QC metrics tested were mitochondrial gene percentage (MT%), number of features ($log_2$(Genes/Features)), and number of molecules ($log_2$(UMI/Counts)). For each dataset, QC metrics were averaged per sample within each overlap category.

**Supplementary Table 1.** Cohen's *d* of QC metrics. Only datasets with more than 3 samples were included in the analysis. QC metrics were averaged per sample within each overlap category. Effect sizes, using Cohen's *d*, were computed between Hidden and other categories. Comparisons with Missed are colored blue-grey. If the metric matches the hypothesis of Hidden being malignant cells -- negligible or small effect sizes with Malignant and moderate or large effect sizes with Not malignant -- it is green. Otherwise, it is red.

| Project | Metric | Malignant | Not malignant | Missed |
|---|---|---|---|---|
| Pancreas (n=5) | % MT | 0.49 (small) | 1.64 (large) | 1.39 (large) |
| | log2(Genes/Features) | -0.55 (moderate) | 2.27 (large) | 1.73 (large) |
| | log2(UMI/Counts) | -0.65 (moderate) | 1.74 (large) | 2.13 (large) |
| Pozniak (n=24) | % MT | -0.72 (moderate) | 0.28 (small) | -0.68 (moderate) |
| | log2(Genes/Features) | -1.42 (large) | 0.37 (small) | -1.12 (large) |
| | log2(UMI/Counts) | -1.21 (large) | 0.55 (moderate) | -0.83 (large) |
| Prostate (n=4) | % MT | -0.09 (negligible) | 0.81 (large) | -0.27 (small) |
| | log2(Genes/Features) | -0.29 (small) | 1.48 (large) | 0.00 (negligible) |
| | log2(UMI/Counts) | -0.35 (small) | 1.44 (large) | -0.02 (negligible) |
| TuPro (n=23) | % MT | -0.41 (small) | 0.12 (negligible) | -0.95 (large) |
| | log2(Genes/Features) | 0.06 (negligible) | 1.31 (large) | 0.73 (moderate) |
| | log2(UMI/Counts) | 0.17 (negligible) | 1.37 (large) | 0.79 (moderate) |
| Uveal melanoma (n=15) | % MT | 0.27 (small) | 0.58 (moderate) | 0.22 (small) |
| | log2(Genes/Features) | -3.68 (large) | -2.77 (large) | -3.04 (large) |
| | log2(UMI/Counts) | -3.40 (large) | -2.27 (large) | -2.68 (large) |

### Supplementary Figure 5

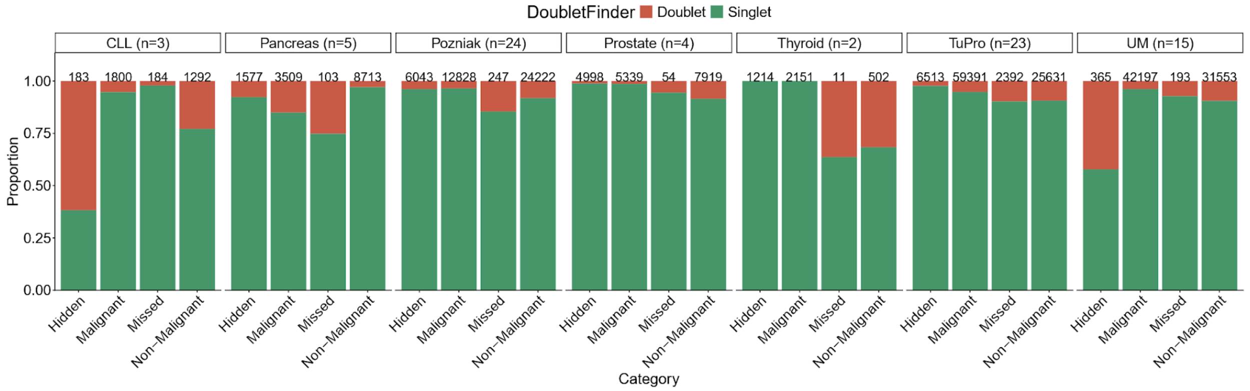


**Supplementary Figure 5.** Doublets and singlets in categories. DoubletFinder was used to identify doublets in samples. Each dataset is grouped by categories and the proportion of doublets and singlets are shown. The number on top of the bar graph indicates the number of cells in the bar below.

**Supplementary Table 2.** Dataset-specific pairwise comparison of DoubletFinder doublet enrichment across Category groups. Within each project, mixed-effects logistic regression (cbind(Doublet, Singlet) ~ Category + (1 | samples)) was used to estimate odds ratios for doublet classification for each Category relative to Hidden. Pairwise contrasts were derived from estimated marginal means. OR < 1 indicates greater doublet enrichment in Hidden, whereas OR > 1 indicates lower doublet enrichment in Hidden. Significant contrasts (Adj. p-value < 0.05) with OR > 1 are shown in green, significant contrasts with OR < 1 are shown in red, and non-significant contrasts are shown in blue-grey. Holm-adjusted p-values and Bonferroni-adjusted confidence intervals are reported. Datasets with fewer than 3 samples were excluded from the analysis. Dataset sample numbers shown.

| Dataset | Contrast | Odds ratio | SE | 95% CI | Adj. p-value |
|---|---|---|---|---|---|
| Pancreas (n=5) | Non-Malignant vs Hidden | 0.104 | 0.017 | 0.07--0.15 | 1.36 x 10^-41 |
| | Malignant vs Hidden | 0.678 | 0.107 | 0.46--0.99 | 0.0281 |
| | Missed vs Hidden | 1.150 | 0.320 | 0.60--2.24 | 0.603 |
| Pozniak (n=24) | Non-Malignant vs Hidden | 2.910 | 0.221 | 2.42--3.49 | 2.15 x 10^-44 |
| | Malignant vs Hidden | 0.889 | 0.074 | 0.73--1.09 | 0.160 |
| | Missed vs Hidden | 4.730 | 0.924 | 2.97--7.55 | 3.37 x 10^-15 |
| Prostate (n=4) | Non-Malignant vs Hidden | 9.390 | 1.350 | 6.65--13.26 | 6.84 x 10^-54 |
| | Malignant vs Hidden | 1.000 | 0.175 | 0.66--1.52 | 0.985 |
| | Missed vs Hidden | 3.800 | 2.330 | 0.88--16.50 | 0.0587 |
| TuPro (n=23) | Non-Malignant vs Hidden | 5.130 | 0.456 | 4.15--6.35 | 2.29 x 10^-75 |
| | Malignant vs Hidden | 2.310 | 0.207 | 1.86--2.87 | 9.34 x 10^-21 |
| | Missed vs Hidden | 4.880 | 0.548 | 3.73--6.39 | 4.71 x 10^-45 |
| UM (n=15) | Non-Malignant vs Hidden | 0.218 | 0.025 | 0.17--0.29 | 1.00 x 10^-41 |
| | Malignant vs Hidden | 0.052 | 0.006 | 0.04--0.07 | 6.06 x 10^-155 |
| | Missed vs Hidden | 0.121 | 0.036 | 0.06--0.25 | 1.33 x 10^-12 |

### Supplementary Figure 6

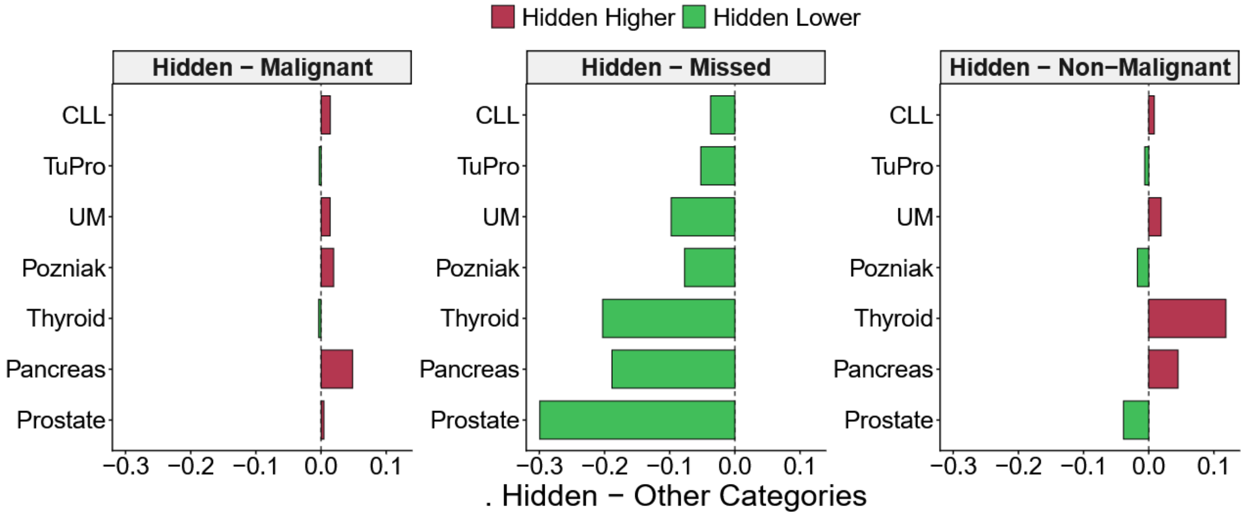


**Supplementary Figure 6.** Difference in hidden ambient RNA contamination compared to other categories. DecontX (28) was run on all samples individually and the median contamination score compared between the different categories for each dataset. Red, positive bars indicate increased median ambient RNA contamination in Hidden and negative, green bars indicate decreased median ambient RNA contamination in hidden.

### Supplementary Figure 7

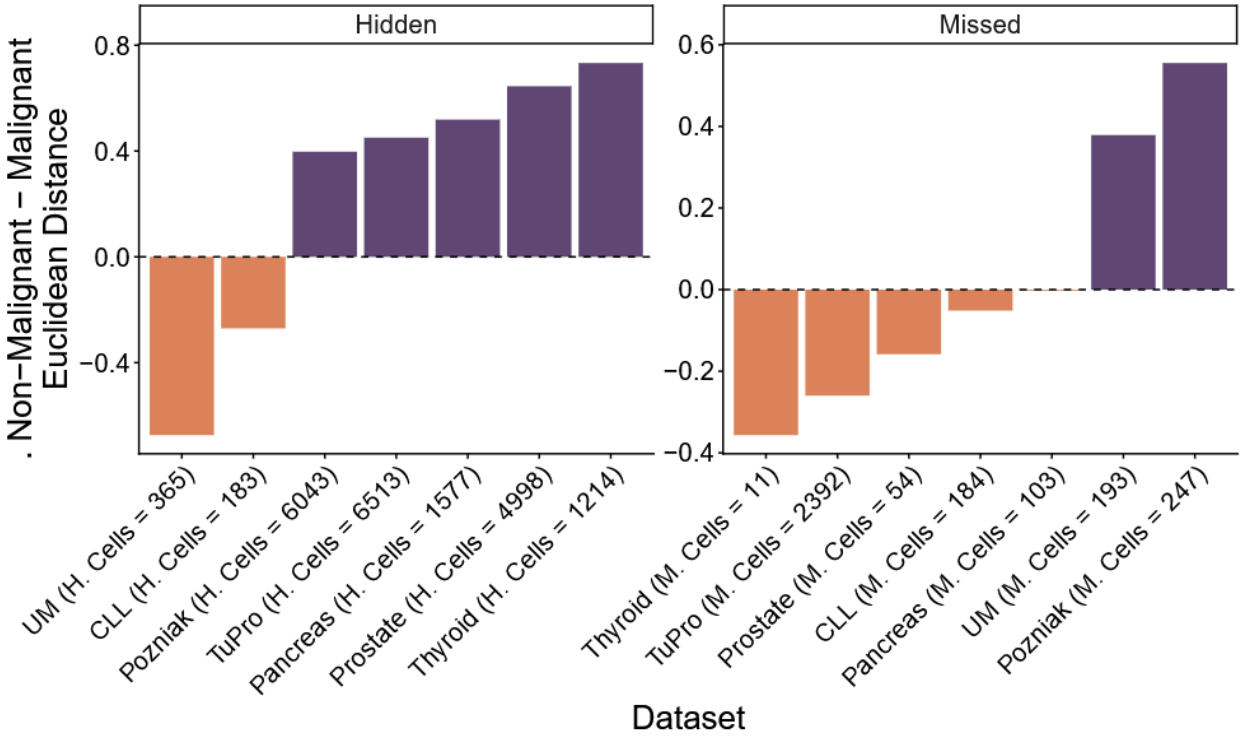

**Supplementary Figure 7.** Euclidean distance of category centroids within datasets. Forty-six Weizmann Institute signatures (4, 41) were applied to the data using UCell (44). Signatures were then selected based on variability across groups assessed using the Kruskal–Wallis $\varepsilon^2$ effect size and interquartile range. Median UCell scores were calculated per category within each dataset. For each dataset, the Euclidean distance between the Hidden or Missed centroid and Non-Malignant or Malignant centroids in signature space was calculated. The difference between Non-Malignant and Malignant to either Hidden or Missed is plotted on the y-axis. Negative distances indicate Hidden or Missed centroids are closer to Non-Malignant centroids, while positive distances indicate Malignant is closer to Hidden or Missed. Numbers of Hidden or Missed cells per cohort are indicated on the x-axis.

### Supplementary Figure 8

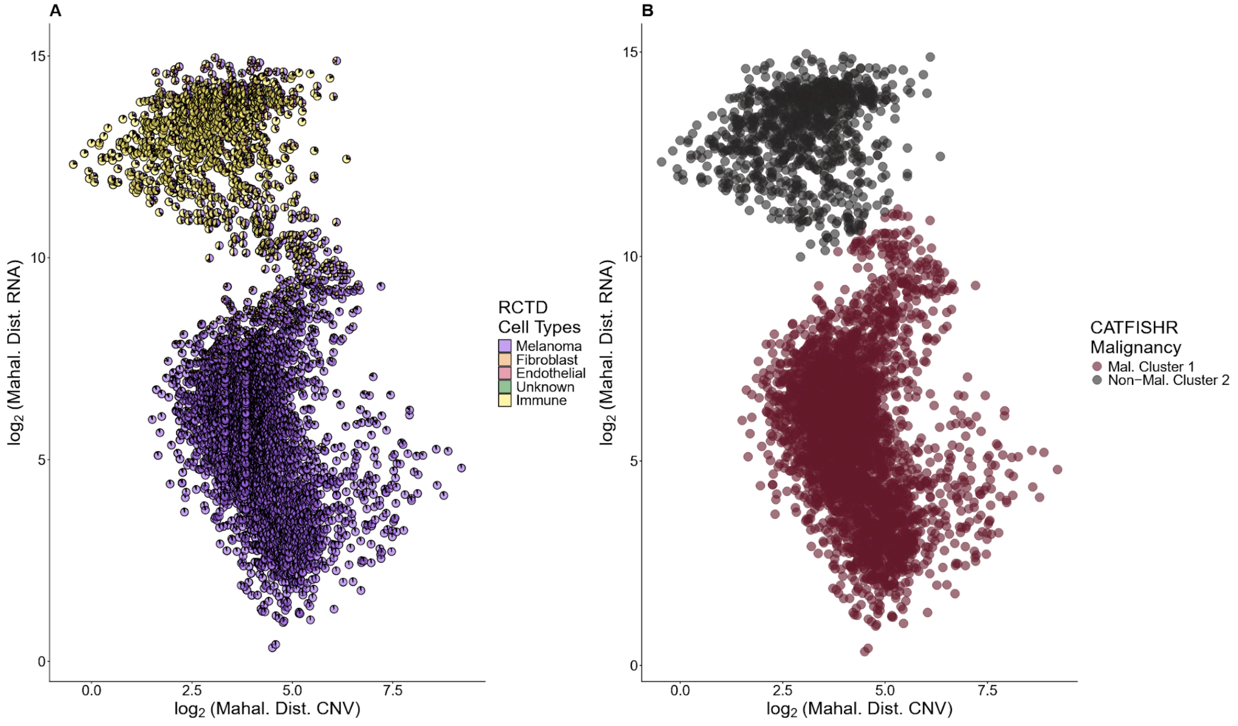


**Supplementary Figure 8.** Visium sample spot RCTD composition and CATFISHR malignancy. A) RCTD cell type composition by spot in Mahalanobis distance space. B) Mean shift clusters and CATFISHR malignancy annotation indicated by color in Mahalanobis distance space.

### Supplementary Figure 9

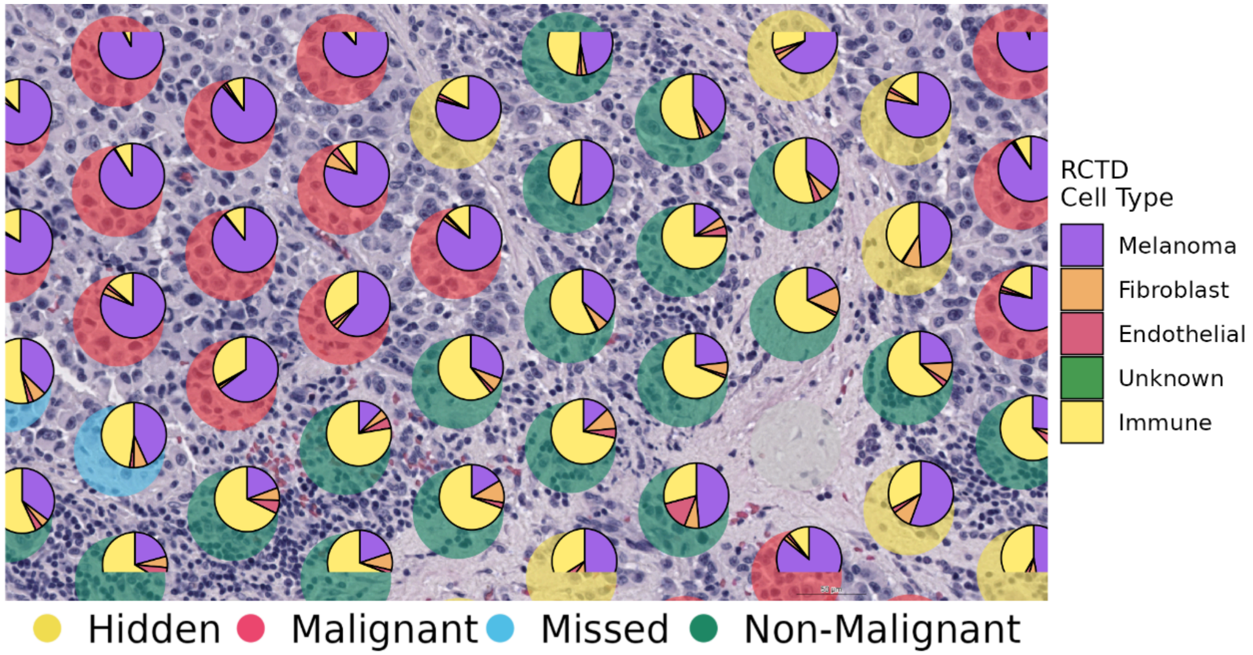


**Supplementary Figure 9.** Visium subsection pie charts. Figure 5F with RCTD cell type (right legend). Category labels visible behind pie charts (bottom legend).

**Supplementary Table 3.** CopyKAT and InferCNV CATFISHR performance comparison summary. The differences in balanced accuracy and F1 score between CATFISHR were calculated per sample. For each comparison, the median, IQR, and comparison outcome are shown. For Δ inferCNV CATFISHR - CopyKAT CATFISHR, runs were considered equivalent ("Equal") if the metric difference was < 0.05. The number of samples meeting this criterion is shown as the numerator, with the comparison-specific number of evaluated samples as the denominator. For the other two comparisons, CATFISHR runs were considered improved relative to the CopyKAT naive algorithm if the metric difference was > 0.05. The number of samples classified as 'Improved' is shown.

**Balanced accuracy**

| | Δ inferCNV CATFISHR – CopyKAT CATFISHR | | | Δ CopyKAT CATFISHR - CopyKAT | | | Δ inferCNV CATFISHR - CopyKAT | | |
|---|---|---|---|---|---|---|---|---|---|
| **Dataset** | **Median** | **IQR** | **Equal** | **Median** | **IQR** | **Improved** | **Median** | **IQR** | **Improved** |
| CLL | -0.0144 | 0.132 | 1/3 | 0.00206 | 0.0855 | 1/3 | 0.0133 | 0.0617 | 1/3 |
| Pancreas | 0.00665 | 0.0852 | 3/5 | -0.00239 | 0.0230 | 0/5 | 0.0199 | 0.178 | 0/5 |
| Prostate | -0.205 | 0.275 | 0/4 | -0.000312 | 0.0382 | 1/5 | -0.186 | 0.356 | 1/4 |
| Thyroid | -0.0841 | 0.0355 | 1/2 | 0.129 | 0.0169 | 2/2 | 0.0444 | 0.0186 | 1/2 |

**F1 score**

| | Δ inferCNV CATFISHR – CopyKAT CATFISHR | | | Δ CopyKAT CATFISHR - CopyKAT | | | Δ inferCNV CATFISHR - CopyKAT | | |
|---|---|---|---|---|---|---|---|---|---|
| **Dataset** | **Median** | **IQR** | **Equal** | **Median** | **IQR** | **Improved** | **Median** | **IQR** | **Improved** |
| CLL | -0.0180 | 0.0727 | 1/3 | 0.00234 | 0.118 | 1/3 | 0.0580 | 0.0823 | 2/3 |
| Pancreas | 0.00798 | 0.244 | 1/5 | -0.00709 | 0.0357 | 0/5 | 0.0137 | 0.293 | 1/5 |
| Prostate | -0.225 | 0.0787 | 0/4 | -0.0000975 | 0.0207 | 1/5 | -0.191 | 0.0887 | 0/4 |
| Thyroid | 0.176 | 0.0695 | 0/2 | 0.145 | 0.0329 | 2/2 | 0.321 | 0.0366 | 2/2 |
